# Primacy coding facilitates effective odor discrimination when receptor sensitivities are tuned

**DOI:** 10.1101/370916

**Authors:** David Zwicker

**Affiliations:** Max Planck Institute for Dynamics and Self-Organization, Am Faßberg 17, 37077 Göttingen, Germany; John A. Paulson School of Engineering and Applied Sciences, Harvard University, Cambridge, MA 02138, USA; Kavli Institute for Bionano Science and Technology, Harvard University, Cambridge, MA 02138, USA

## Abstract

The olfactory system faces the difficult task of identifying an enormous variety of odors independent of their intensity. Primacy coding, where the odor identity is encoded by the receptor types that respond earliest, is one possible representation that can facilitate this task. So far, it is unclear whether primacy coding facilitates typical olfactory tasks and what constraints it implies for the olfactory system. In this paper, we develop a simple model of primacy coding, which we simulate numerically and analyze using a statistical description. We show that the encoded information depends strongly on the number of receptor types included in the primacy representation, but only weakly on the size of the receptor repertoire. The representation is independent of the odor intensity and the transmitted information is useful to perform typical olfactory tasks, like detecting a target odor or discriminating similar mixtures, with close to experimentally measured performance. Interestingly, we find situations in which a smaller receptor repertoire is advantageous for identifying a target odor. The model also suggests that overly sensitive receptor types could dominate the entire response and make the whole array useless, which allows us to predict how receptor arrays need to adapt to stay useful during environmental changes. By quantifying the information transmitted using primacy coding, we can thus connect microscopic characteristics of the olfactory system to its overall performance.

**Author summary:** Humans can identify odors independent of their intensity. Experimental data suggest that this is accomplished by representing the odor identity by the earliest responding receptor types. Using theoretical modeling, we here show that such a primacy code allows discriminating odors with close to experimentally measured performance. This performance depends strongly on the number of receptors considered in the primacy code, but the receptor repertoire size is less important. The model also suggests a strong evolutionary pressure on the receptor sensitivities, which could explain observed receptor copy number adaptations. Taken together, the model connects detailed molecular measurements to large-scale psycho-physical measurements, which will contribute to our understanding of the olfactory system.

## Introduction

The olfactory system identifies and discriminates odors for solving vital tasks like navigating the environment, identifying food, and engaging in social interactions. These tasks are complicated by the enormous variety of odors, which vary in composition and in the concentrations of their individual molecules. In particular, the olfactory system needs to separately recognize the odor identity (what is there?) and the odor intensity (how much is there?). For instance, the identity is required to decide whether to approach or avoid an odor source, whereas the intensity information is important for localizing it. It is unclear how these two odor properties are separated.

Odors are sensed by olfactory receptors that have distinct responses to different odor molecules. Generally, each receptor responds to a wide range of odors and each odor activates many receptor types. The resulting combinatorial code allows to distinguish odor identities [1–3], but also depends on the odor intensity, since receptors respond stronger to more concentrated molecules [4]. To obtain an intensity-invariant code in the olfactory cortex [5, 6], the neural information is processed in the olfactory bulb in mammals and the antenna lobe in insects [7–9]. For instance, inhibiting neurons in the olfactory bulb [10, 11] affect the neurons processing the receptor activities globally [12–18], which could result in a concentration-invariant representation of the odor identity [19, 20]. However, we showed that such a normalized representation still depends strongly on the number of ligands in a mixture and might thus not be optimal for solving olfactory tasks [21]. An alternative to these normalized representations is rank coding, where the order in which the receptors are excited is used to encode the odor identity robustly and independently of the odor intensity [22]. Indeed, experiments suggests that odors are encoded robustly by the receptor types that respond within a given time window after sniff onset [23, 24]. In particular, the odor identity could be robustly encoded by a fixed number of the receptors that respond first, which is known as primacy coding [23, 25]. So far, it is unclear whether this simple coding scheme is sufficient to explain the remarkable discriminatory capability of the olfactory system.

In this paper, we consider a simple model of primacy coding and investigate how well it represents the identity of complex odors. In particular, we identify how much information is transmitted and how well this information can be used to perform typical olfactory tasks, like identifying a target odor in a background or discriminating odor mixtures. Our model thus links parameters of the primacy code with results from typical psychophysical experiments. We show that primacy coding provides a robust and compact representation of the odor identity over a wide range of odors, independent of the odor intensity. However, this good performance of the olfactory system hinges on tuned receptor sensitivities, which suggests that there is a strong selective pressure to adjust the sensitivities on evolutionary and shorter timescales.

## Results

We describe odors by concentration vectors ***c*** = (*c*_1_, *c*_2_, …,*c_N_L__*), which determine the concentrations *c_i_ >* 0 of all detectable ligands to the olfactory receptors. The number *N*_L_ of possible ligands is at least *N*_L_ = 2300 [26] although the realistic number is likely much larger [27]. Typical odors contain only tens to hundreds of ligands, implying that most *c_i_* are zero. The statistics of natural odors are difficult to measure [28]. We thus consider a broad class of odor distributions, where each ligand *i* has a probability *p_i_* to appear in an odor. For simplicity, we neglect correlations in their appearance, so the mean number *s* of ligands in an odor is *s* = ∑*_i_ p_i_*. To model the broad distribution of ligand concentrations, we choose the concentration *c_i_* of ligand *i* from a log-normal distribution with mean *μ_i_* and standard deviation *σ_i_* if the ligand is present. Consequently, the mean concentration of a ligand in any odor reads ⟨*c_i_* ⟩= *p_i_μ_i_* and the associated variance is 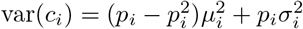. For simplicity, we consider ligands with equal statistics in this paper, so the distribution *P*_env_(***c***) of odors is characterized by the three parameters *p_i_* = *p*, *μ_i_* = *μ*, and *σ_i_* = *σ*.

### Simple model of primacy coding

Odors are detected by an array of receptors in the nasal cavity in mammals and on the antenna in insects. The receptor array consists of *N*_R_ different receptor types, which each are expressed many times. Typical numbers are *N*_R_ ≈ 50 in flies [7], *N*_R_ ≈ 300 in humans [29], and *N*_R_ ≈ 1000 in mice [30]. The excitations of all receptors of the same type are accumulated in an associated glomerulus in the olfactory bulb in mammals and the antennal lobe in insects [31]. Since this convergence of the neural information mainly improves the signal-to-noise ratio, we here capture the excitation of the receptors on the level of glomeruli; see Fig. 1A. The excitation *e_n_* of glomerulus *n* can be approximated by a linear map of the odor ***c*** [4, 32],

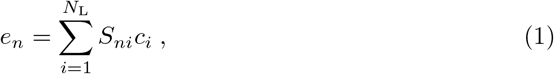

where *S_ni_* denotes the effective sensitivity of glomerulus *n* to ligand *i*. Note that *S_ni_* is proportional to the copy number of receptor type *n* if the response from all individual receptors is summed [33].

**Fig 1.**
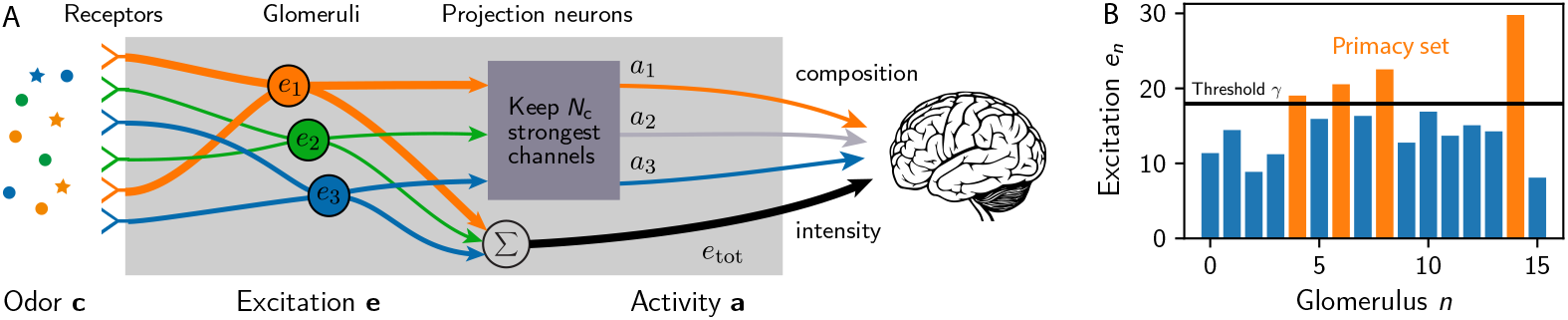
Simple model of primacy coding. (A) Schematic picture of the signal processing in the olfactory bulb: An odor comprised of many ligands excites the olfactory receptors and the signals from all receptors of the same type are accumulated in respective glomeruli. The glomeruli with the strongest (earliest) excitations encode the odor composition, whereas the odor intensity could be encoded separately. (B) Excitations of *N*_R_ = 16 glomeruli for an arbitrary odor. The *N*_C_ = 4 glomeruli with the highest excitations, above the threshold *γ*, form the primacy set (orange bars).

The sensitivity matrix *S_ni_* could in principle be determined by measuring the response of each glomerulus to each possible ligand. However, because the numbers of receptor and ligand types are large, this is challenging and only parts of the sensitivity matrix have been measured, e. g., in humans [34] and flies [35]. We showed that the measured matrix elements are well described by a log-normal distribution with a standard deviation *λ* ≈ 1 of the underlying normal distribution [33]. Motivated by these observations, we here consider random sensitivity matrices, where each element *S_ni_* is chosen independently from the same log-normal distribution, which is parameterized by its mean ⟨*S_ni_*⟩ = *S*̄ and variance 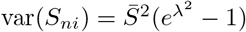. Since these receptor sensitivities are broadly distributed, they might not include specific receptors related to innate behavior [36], but they can collectively discriminate concentration differences of several orders of magnitude [33].

The odor representation on the level of glomeruli excitations *e_n_* depends strongly on the odor intensity *c*_tot_ = ∑*i c_i_*, which complicates the extraction of the odor identity determined by the relative concentrations. A concentration-invariant representation could be achieved by normalizing the excitations by the mean excitation [16], which leads to an efficient neural representation on the level of projection neurons [21]. However, recent experimental data suggest an alternative encoding based on the timing of the glomeruli excitation [23]. The key idea of this primacy coding is that the set of receptor types that are excited first is independent of the total concentration *c*_tot_ and thus provides a concentration-invariant representation. In the simple situation where bound ligands only affect the strength of the receptor output, but not the signaling dynamics, the receptors that first cross a threshold are the ones with the largest excitation. For simplicity, we also neglect the order in which excitations cross the threshold, in contrast to rank coding. Taken together, the primacy code is then given by the identity of the *N*_C_ glomeruli with the largest excitation, which is known as the primacy set [37].

The primacy set can be represented by a binary vector 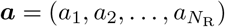, where *a_n_* = 1 implies that glomerulus *n* belongs to the primacy set and is active, while *a_n_* = 0 denotes an inactive glomerulus not belonging to the primacy set. Since the active glomeruli have the highest excitation, they can be identified using an excitation threshold *γ*; see Fig. 1B. Consequently, the activities are given by

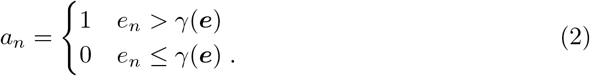

Physiologically, the activities *a_n_* could be encoded by projection neurons in insects and mitral and tufted cells in mammals. These neurons receive excitatory input from one glomerulus [38] and are inhibited by a local network of granule cells [20, 31]. These granule cells basically integrate the activity of all glomeruli [39] and could inhibit the glomeruli once a threshold is reached. Taken together, this would imply primacy coding since only the glomeruli that respond earliest would be activated. For simplicity, we consider the case where the number *N*_C_ of active glomeruli is fixed and does not depend on the odor ***c***. The associated constraint

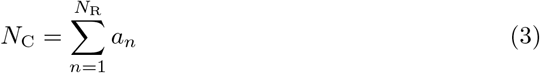

determines the threshold *γ*. The activity pattern ***a*** is sparse since only a fraction *N*_C_*/N*_R_ of all glomeruli are activated. Moreover, ***a*** is concentration-invariant, since the odor intensity *c*_tot_ does not affect ***a***. This is because multiplying the concentration vector ***c*** by a constant factor changes both the excitations *e_n_* and the threshold *γ* by the same factor, so that ***a*** given by Eq. (2) is unaffected. In essence, only relative excitations are relevant for our model of primacy coding.

The amount of information that can be learned about the odor ***c*** by observing the activity pattern ***a*** is quantified by the mutual information *I* given by

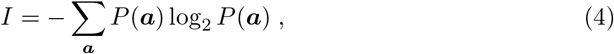

where the probability *P* (*a*) of observing an output ***a*** depends on the odor environment *P*_env_(***c***) as well as the properties of the olfactory system, which in our model are quantified by *N*_C_, *N*_R_, and *λ*.

In an optimal receptor array, each output ***a*** occurs with equal probability when encountering odors distributed according to *P*_env_(***c***) [33]. In the case of primacy coding, only outputs with exactly *N*_C_ active receptor types are permissible. Consequently, in the optimal representation each receptor type would be activated with a probability ⟨*a_n_*⟩ = *N*_C_*/N*_R_ and all types would be uncorrelated, cov(*a_n_, a_m_*) = 0 for *n* ≠ *m*. The associated information

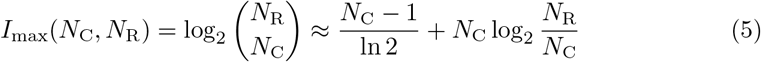

provides an upper bound for *I* given by Eq. (4). Here, the approximation on the right hand side is obtained using Stirling’s formula for large receptor repertoires (*N*_R_ ≫ *N*_C_). Note that primacy coding contains much less information than simple binary coding (where all glomeruli are considered [33, 40]) and rank coding (where the order of activation of the first *N*_C_ glomeruli is also included [22]); see Fig. 2A. Nonetheless, we will show below that primacy coding provides useful information for solving typical olfactory tasks and can even outperform alternatives encoding more information.

**Fig 2.**
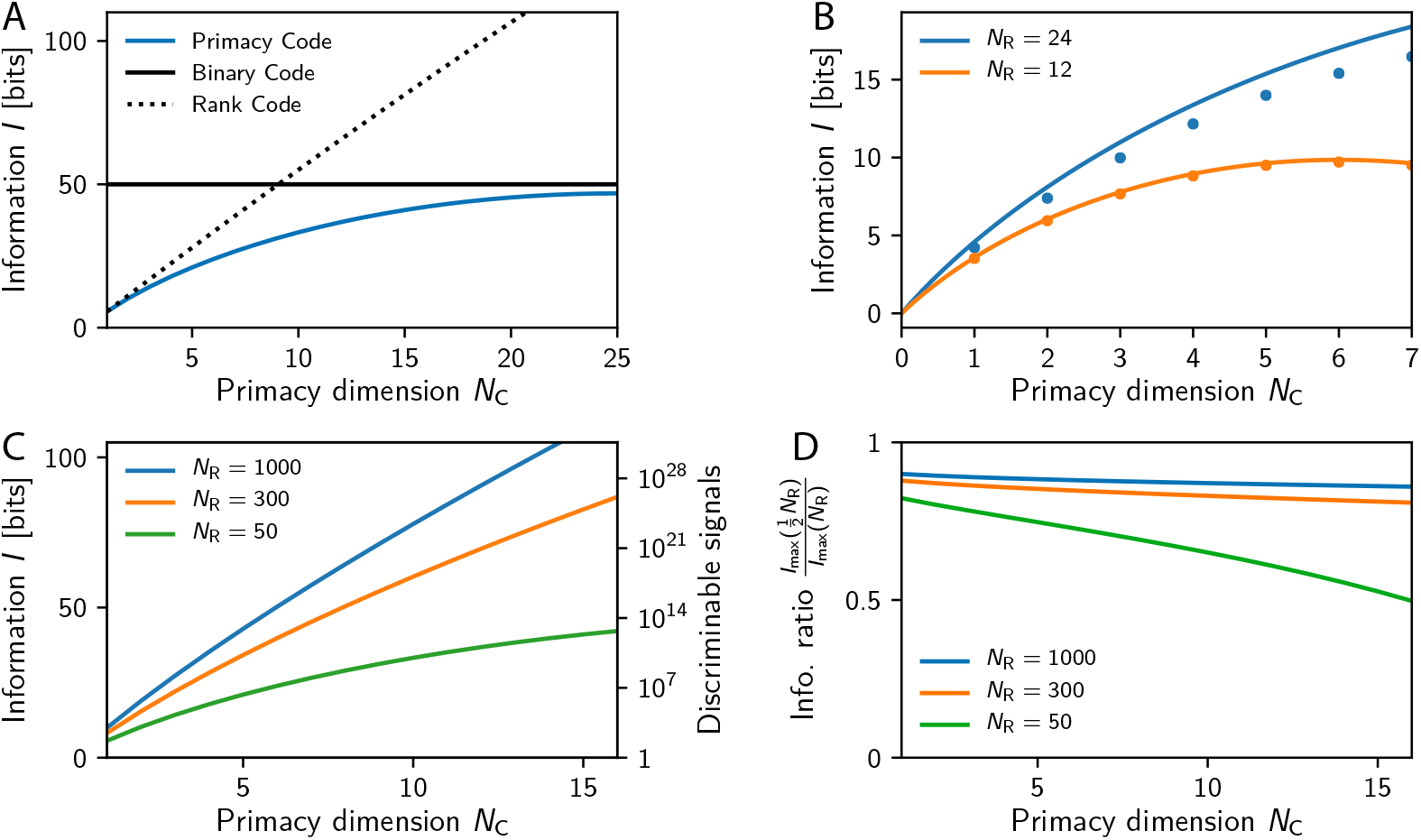
Transmitted information *I* increases strongly with primacy dimension *N*_C_ and weakly with receptor repertoire size *N*_R_. (A) Comparison of the transmitted information *I* in primacy coding, binary coding, and rank coding for *N*_R_ = 50. (B) The maximally transmitted information *I*_max_ (solid lines) given by Eq. (5) is compared to numerical estimates of *I* (dots; *n* = 10^7^, error smaller than symbol size) obtained from ensemble averages of Eq. (4). Model parameters are *N*_L_ = 512, *μ* = *σ* = 1, *s* = 16, and *λ* = 1, implying *ζ* ≈ 0.1. (C) *I*_max_ as a function of *N*_C_ for several *N*_R_. The right axis indicates the maximal number 2^*I*^ of distinguishable signals. (D) Reduction of *I*_max_ when half the receptor types are removed as a function of *N*_C_ for various *N*_R_.

### Transmitted information depends weakly on receptor repertoire

We start by analyzing the information *I* transmitted by the primacy code using numerical ensemble averages of Eqs. (1)–(4); see Methods and Models. Fig. 2B shows that *I* is very close to the maximal information *I*_max_ given by Eq. (5), which is obtained when all receptor types have equal activity and are uncorrelated [33]. This indicates that the primacy code uses the different receptor types with similar frequency and that correlations between them are negligible. The expression for *I*_max_ implies that the information grows linearly with the primacy dimension *N*_C_, but only logarithmically with the number *N*_R_ of receptor types. Consequently, the number of distinguishable signals, given by 2^*I*^, grows strongly with *N*_C_, but the dependence on the repertoire size is weaker. Given equal *N*_C_, our model thus predicts that the transmitted information in mice is only twice that of flies, although mice possess about 20 times as many receptor types; see Fig. 2C. However, the number of discriminable signals changes by many orders of magnitudes because of the exponential scaling with *I*.

The logarithmic scaling of the transmitted information *I* with the receptor repertoire size *N*_R_ could explain why the ability of rats to discriminate odors is not significantly affected when half the olfactory bulb is removed in lesion experiments [41, 42]. If this operation removes half the receptor types, our model implies that the transmitted information *I* is lowered by *N*_C_ bits; see Eq. (5). This corresponds to a reduction of *I* by about 10 % in rats where *N*_R_ ≈ 1000; see Fig. 2D. Conversely, the transmitted information can be reduced by almost 50 % in flies, which have a much smaller receptor repertoire of *N*_R_ ≈ 50. Our model thus predicts that lesion experiments have a much more severe affect on the performance of animals with smaller receptor repertoires.

Taken together, this first analysis already suggests that the primacy code provides a robust odor representation, which is sparse, concentration-invariant, and depends only weakly on the details of the receptor array. However, for this representation to be useful to the animal, it needs to allow solving typical olfactory tasks.

### Primacy coding discriminates odors efficiently

Typical olfactory tasks involve detecting a ligand in a background, detecting the addition of a ligand to a mixture, and discriminating similar mixtures. All these tasks involve discriminating odors with common ligands, implying that the associated primacy sets are correlated. In particular, discriminating similar odors will be impossible if their primacy sets are identical. To see when discrimination is possible, we quantify the distance *d* between two primacy sets by simply counting the number of glomeruli with different activities.

#### Discriminating uncorrelated odors

To build an intuition for the distance *d* between primacy sets, we start by considering two uncorrelated odors. In this case, each receptor type has an expected activity of ⟨*a_n_*⟩ = *N*_C_*/N*_R_ and the resulting distance reads *d_∗_* = 2*N*_C_(1 *− N*_C_*N*_R_^−1^), which implies that two uncorrelated odors will typically be distinguishable (⟨*d*⟩ *≥* 2), even for very small primacy dimension *N*_C_. Moreover, this expression implies that odors become more easily discriminable when *N*_C_ is increased, whereas increasing the receptor repertoire size *N*_R_ has a negligible effect in the typical case *N*_C_ ≪ *N*_R_. This is similar to the scaling of the transmitted information *I* discussed above.

#### Detecting the presence of a target odor in a background

One simple task where correlated primacy sets matter is the detection of a target odor in a distracting background. To understand when a target can be detected, we analyze how the primacy set ***a*** changes when a single ligand at concentration *c*_t_ is added to a background ligand at concentration *c*_b_. Because of concentration-invariance, only the relative target concentration *c*_t_*/c*_b_ matters and we expect that the target is easier to detect when it is more concentrated (larger *c*_t_*/c*_b_). Fig. 3A shows that this is indeed the case, since the mean change ⟨*d*⟩ in the primacy set ***a*** increase with *c*_t_*/c*_b_. Moreover, ⟨*d*⟩ increases with the primacy dimension *N*_C_ in the same way as the distance *d_∗_* of uncorrelated odors (see inset). In fact, ⟨*d*⟩ must approach *d_∗_* when the target dominates the background (*c*_t_*/c*_b_ *→ ∞*). This scaling implies that the receptor repertoire size *N*_R_ only has a weak effect on ⟨*d*⟩, which is confirmed by Fig. 3B. Surprisingly, the dependence on *N*_R_ is not monotonic and very dilute odors (small *c*_t_*/c*_b_) are actually more difficult to discriminate with larger receptor repertoires.

**Fig 3.**
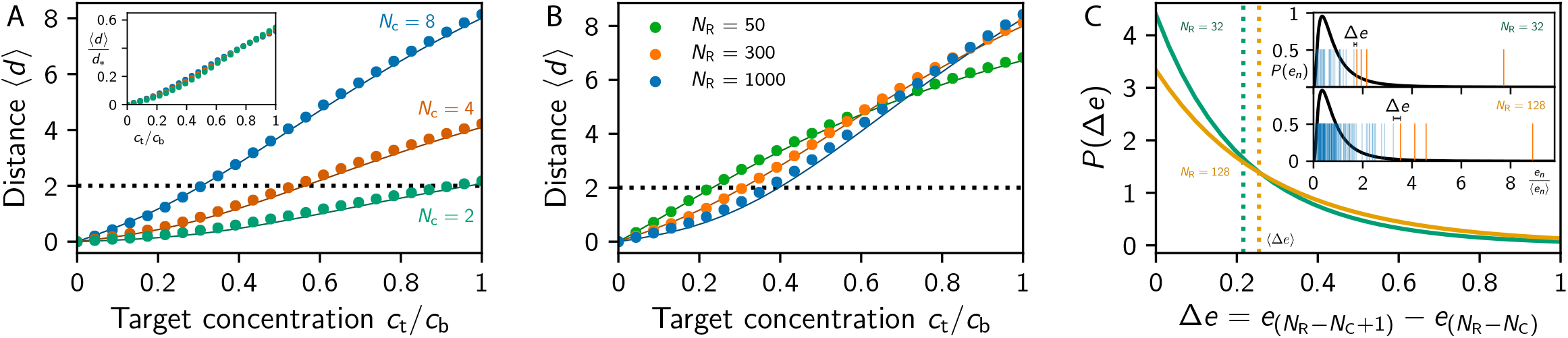
Detection of a target ligand in a background. (A-B) Mean change ⟨*d*⟩ in the primacy set when the target at concentration *c*_t_ is added to the background ligand at concentration *c*_b_ as a function of *c*_t_*/c*_b_ for (A) various *N*_C_ at *N*_R_ = 300 (inset: same data rescaled by *d_∗_*) and (B) various *N*_R_ at *N*_C_ = 8. Numerical simulations (dots; sample size: 10^5^) are compared to the theoretical prediction (lines) obtained using the statistical model; see Methods and Models. The dotted line indicates the discrimination threshold ⟨*d*⟩ = 2. (C) Distribution of the difference ∆*e* between the excitations just above and below the threshold for *N*_R_ = 32 (green line) and *N*_R_ = 128 (orange line); see Methods and Models. The dotted vertical lines indicate the mean ⟨∆*e*⟩, obtained from Eq. (12). The inset shows realizations of *N*_R_ = 32 (upper panel) and *N*_R_ = 128 (lower panel) excitations (vertical bars) drawn from the same excitation distribution (black lines). The orange bars indicate the primacy set consisting of the *N*_C_ = 4 largest excitations. (A–C) Remaining parameters are given in Fig. 2B.

The fact that increasing the receptor repertoire size *N*_R_ can impede the detection of the target odor can be understood in a simplified statistical model, where we calculate the expected distance ⟨*d*⟩ using ensemble averages over sensitivity matrices; see Methods and Models. Since the primacy set ***a*** corresponds to the *N*_C_ receptor types with the largest excitations, ***a*** will only change when adding the target odor brings the excitation of an inactive receptor type above the excitation of a previously active one. Intuitively, this is more likely when the difference ∆*e* between the excitation of the weakest active receptor type and the strongest inactive one is small. Fig. 3C shows that large ∆*e* are more likely for larger *N*_R_, essentially because the distribution of the glomeruli excitation *e_n_* has a heavy tail, so that sampling more excitations leads to larger gaps between the largest excitations. In this case, it is less likely that perturbing the odor changes the order of the excitations and thus the primacy set. Consequently, the maximal concentration *c*_t_*/c*_b_ at which a target can still be detected increases with the receptor repertoire size *N*_R_; see Fig. 3B. In contrast, increasing the primacy dimension *N*_C_ always improves the detection limit.

#### Detecting the addition of a ligand to a mixture

So far, we considered simple odors consisting of single ligands. However, realistic odors are comprised of many different ligands and a more realistic olfactory task is thus the detection of a target in a background of many distracting ligands. For simplicity, we first consider the case where all ligands have the same concentration and we only vary the number *s* of ligands in the background odor. Using ensemble averages over sensitivity matrices, we show in Fig. 4A that the discriminability ⟨*d*⟩ decreases both with larger mixture sizes *s* and smaller primacy dimension *N*_C_. In experiments, humans can detect the presence or absence of a ligand for mixtures of up to about 16 ligands [43] and mice perform even better [44]. To see whether this performance is achievable with primacy coding, we use our statistical model to determine the maximal mixture size *s_∗_* at which the addition of the target odor can still be detected (i. e. when ⟨*d*⟩ ≥ 2). The solid lines in Fig. 4B show that *s_∗_* = 16 is feasible for *N*_C_ ≈ 7 in humans if all ligands have the same concentration 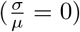. However, if the concentration of the individual ligands is drawn from a distribution with significant variance, a much larger primacy dimension of *N*_C_ ≈ 15 is necessary to still detect the absence or presence of an additional ligand for *s* = 16 (dashed lines in Fig. 4B, 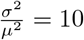).

**Fig 4.**
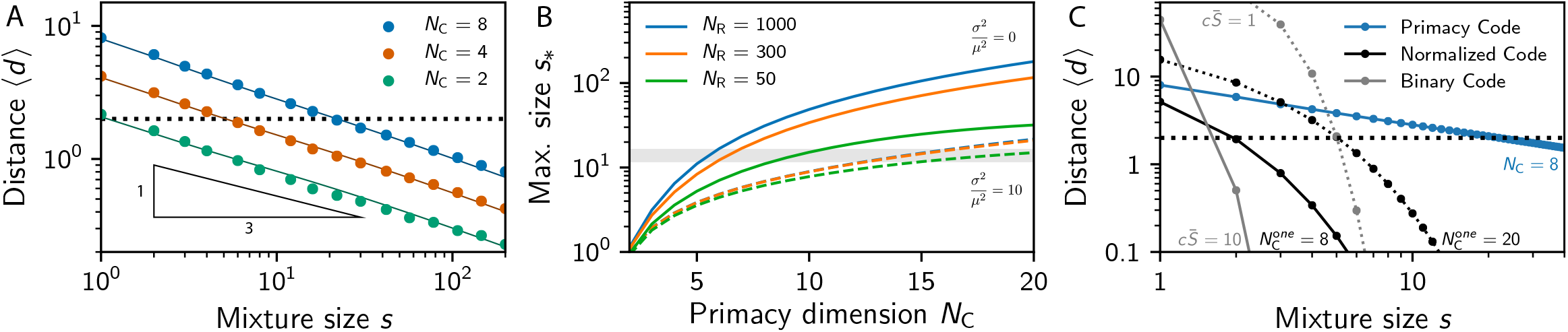
Detection of the presence of a ligand in a mixture. (A) Mean change ⟨*d*⟩ in the primacy set when a ligand is added to a mixture with *s* ligands for various *N*_C_ and *N*_R_ = 300. (B) Maximal mixture size *s_∗_* at which adding a ligand results in *d* = 2 as a function of *N*_C_ for various *N*_R_. Shown is the case of uniform ligand concentrations (solid lines; *σ* = 0) and distributed concentrations (dashed lines, *σ*^2^*/μ*^2^ = 10). The gray band indicates the maximal mixture sizes humans can resolve [43]. (C) Comparison of the primacy code (blue; *N*_C_ = 8) to a normalized code (black) and a binary code (gray) for various *s*. In the normalized code, glomeruli are active when their excitation exceeds *α* times the mean excitation [21]. Here, *α* is determined by the indicated number 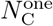 of glomeruli activated by a single ligand. In the binary code, glomeruli are active when their excitation exceeds the fixed threshold, so the activity depends on the odor intensity, explaining the strong dependence on the ligand concentration *c* [33]. In all models, we considered log-normally distributed *S_ni_* with var(*S_ni_*) = 1.72 *S̄*^2^ corresponding to *λ* = 1 and fixed ligands concentrations (*σ* = 0). (A,B) Remaining parameters are given in Fig. 2B. (A,C) The dotted line indicates the discrimination threshold ⟨*d*⟩ = 2.

Interestingly, we find that target odors can be detected more reliably when the background at a given total concentration *c*_b_ consists of more ligands. This can be seen by comparing single-ligand backgrounds (Fig. 3A) with multi-ligand backgrounds (Fig. 4A), where the effective target concentration is *c*_t_*/c*_b_ = 1*/s*. Considering *N*_C_ = 8, the target can only be detected for *c*_t_*/c*_b_ ≲ 1*/*3 in the single-ligand case, while the ratio can be much smaller (*c*_t_*/c*_b_ ≲ 1*/*20) for multiple ligands. This puzzling result can again be understood in the simplified statistical model, which predicts that the variance of the excitations associated with the background odor is smaller if this odor is comprised of many ligands; see Eq. (8) in Methods and Models. This smaller variance implies smaller ∆*e*, so that adding the target has a higher chance of shuffling the order of the excitations to change the primacy set. The same logic implies that the target is easier to detect when the concentrations of the background ligands vary less, which is confirmed by Fig. 4B. Taken together, numerical results and the statistical model suggest that a target odor is easier to notice if the background odor contains many ligands and small concentration variations.

Primacy coding permits the detection of the addition of ligands to mixtures more efficiently than alternative simple encodings. To show this, we also calculate the mean change ⟨*d*⟩ of the activity when a ligand is added to a mixture in two alternative models that have been discussed in the literature; see Fig. 4C. First, we consider a normalized code where glomeruli are active when their excitation normalized to the mean excitations exceeds a threshold value *α*. We showed in [21] that the encoded information and the discriminability strongly depends on the mixtures size *s* in this model. Consequently, a normalized code cannot detect the addition of a ligand at large *s* while at the same time providing a sparse response for individual ligands (small *N*_C_*/N*_R_); see Fig. 4C. In an even simpler model of the olfactory system, glomeruli do not interact at all and are simply activated when their excitation exceeds a threshold [40]. This binary code is not sparse and is strongly affected by the odor intensity, implying that mixtures cannot be discriminated over any significant concentration range [33]. Fig. 4C shows that the discriminability measured by ⟨*d*⟩ decreases much more slowly with the mixture size *s* in primacy coding compared to the alternatives. Taken together, primacy coding provides odor discriminability on physiologically relevant levels using a sparse code for all mixtures sizes.

#### Discriminating similar mixtures

To consider the discrimination of similar odors that have common ligands, we next consider odors that each contain *s* ligands, sharing *s*_B_ of them. Such odors are uncorrelated (⟨*d*⟩ = *d_∗_*) when they do not share any ligands (*s*_B_ = 0) and they are identical (⟨*d*⟩ = 0) when they share all ligands (*s*_B_ = *s*). Between these two extremes, the expected distance ⟨*d*⟩ of the primacy sets of the two odors can be determined by a numerical ensemble average over sensitivities and by the statistical model; see Methods and Models. Fig. 5 shows that both methods predict that more similar odors are harder to discriminate. However, the discriminability of odors only depend on their relative similarly (the fraction of shared ligands) and is independent of the total number of ligands in the odor, consistent with psychophysical experiments [45]. Our model predicts that odors should be distinguishable even if they differ by only about 10 % for *N*_C_ ≳ 4.

**Fig 5.**
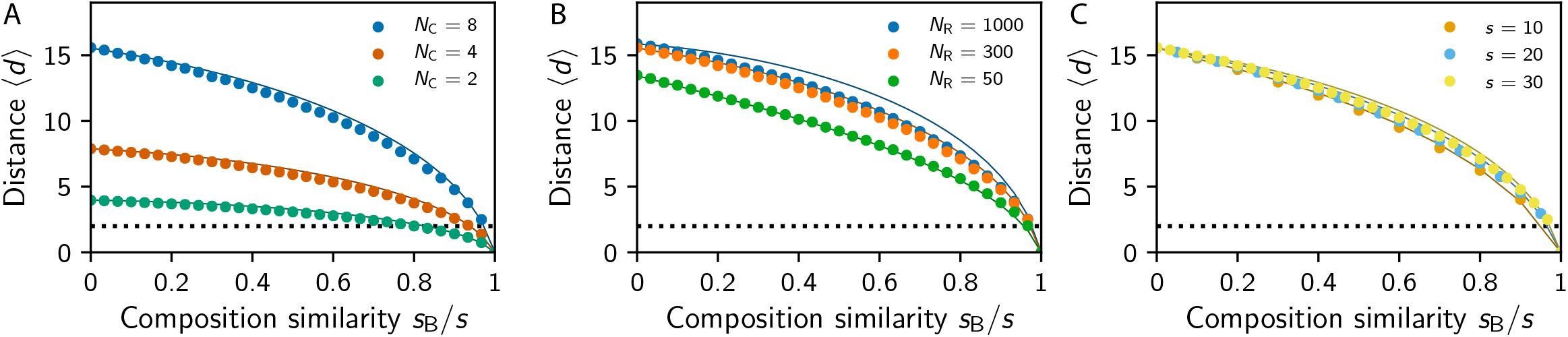
More similar odor mixtures are more difficult to discriminate. Shown is the mean distance ⟨*d*⟩ between the primacy sets of two mixtures with *s* ligands each, sharing *s*_B_ of them. (A) Distance ⟨*d*⟩ for various *N*_C_ at *N*_R_ = 300 and *s* = 30. (B) Distance ⟨*d*⟩ for various *N*_R_ at *N*_C_ = 8 and *s* = 30. (C) Distance *(d)* for various mixture sizes *s* at *N*_R_ = 300, *N*_C_ = 8. (A–C) The dotted line indicates the discrimination threshold ⟨*d*⟩ = 2. Remaining parameters are given in Fig. 2B.

### Overly sensitive receptors degrade the coding efficiency

So far, we calculated the transmitted information and tested the discrimination performance of primacy coding under the assumption that all receptor types behave similarly. In fact, we established that the maximal information is achieved when all receptor types are activated with equal probability *N*_C_*/N*_R_. However, neither the receptor sensitivities nor the odors themselves are distributed equally in realistic situations. Variations in these quantities affect the transmitted information and thus the usefulness of the primacy code. For instance, the transmitted information decreases if a single receptor is activated less often than all the others; see Fig. 6A. This effect is small, since in the worst case the receptor is never active and the transmitted information thus corresponds to an array with the receptor removed. Conversely, having a receptor that is active more often than all others can have a much more severe effect; see Fig. 6A. In fact, if the receptor type is more than three times as active, the transmitted information *I* is lower than if the receptor type was remove completely; see Methods and Models. This indicates that receptors can shadow the response of other receptors and thus be detrimental to the overall array when they are overly sensitive.

**Fig 6.**
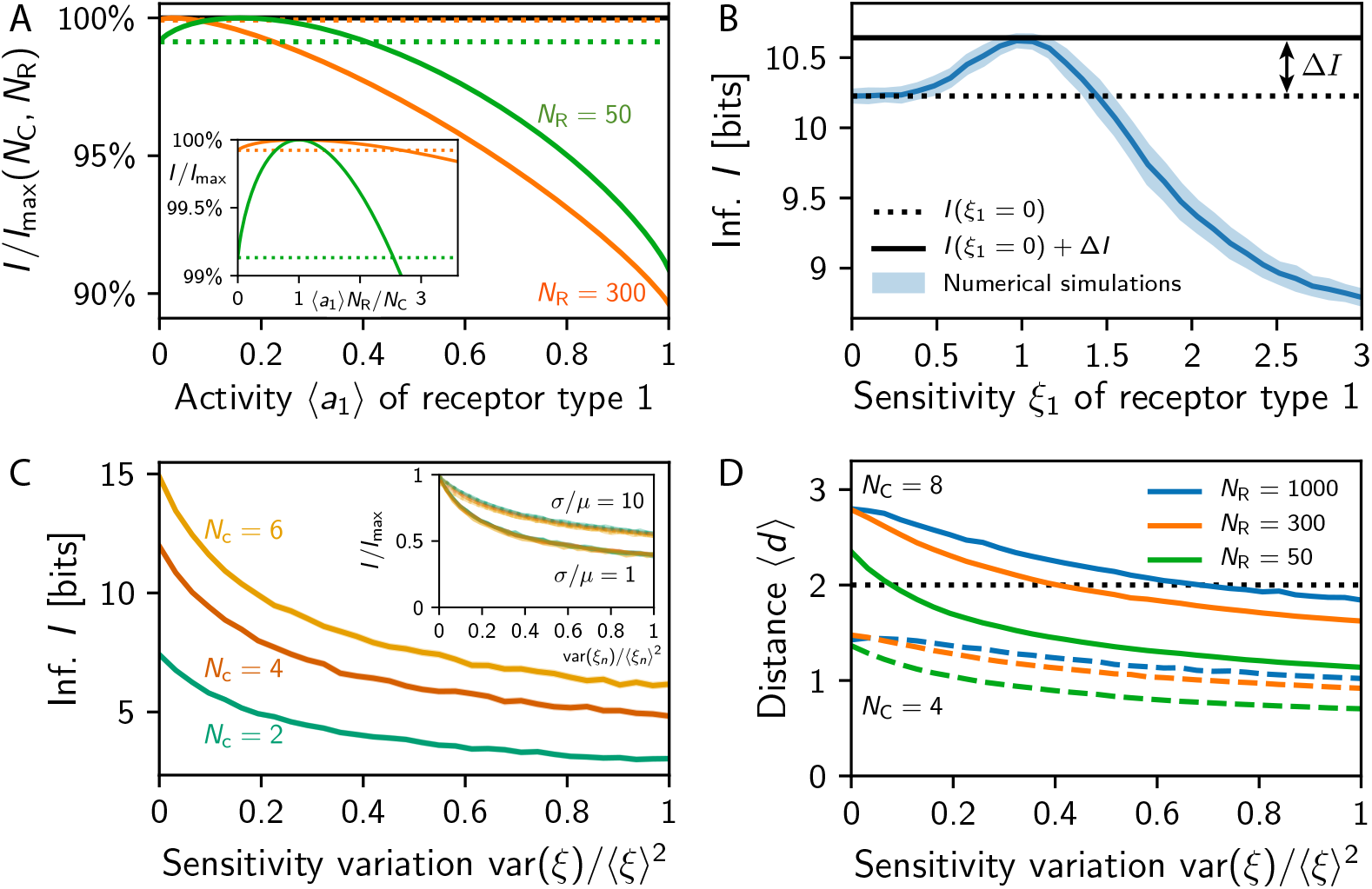
Variations in receptor activities deteriorate the array performance. (A) Information *I* relative to *I*_max_ given in Eq. (5) as a function of the mean activity ⟨*a*_1_⟩ of the first receptor type while all others are unchanged for *N*_C_ = 8. Dotted lines indicate *I*_max_(*N*_C_*, N*_R_ − 1). Inset: Same data for the activity rescaled by *N*_C_*/N*_R_. (B) Numerical simulation of *I* as a function of the sensitivity factor *ξ*_1_ of the first receptor type for *N*_R_ = 16, *N*_C_ = 4, and *ξ_n_* = 1 for *n ≥* 2. (C) *I* for log-normally distributed sensitivity factors *ξ_n_* as a function of the distribution width var(*ξ*)*/* ⟨*ξ*⟩^2^ for various *N*_C_ at *N*_R_ = 20. The inset shows that the scaled information *I/I*_max_ collapses as a function of *N*_R_ = 10, 20 and *N*_C_ = 2, 4, 6 for different widths *σ/μ* of the concentration distribution. (D) Mean change ⟨*d*⟩ in the primacy set caused by adding a ligand to a mixture as a function of var(*ξ*)*/* ⟨*ξ*⟩^2^ for various *N*_R_ and *N*_C_. The dotted black line indicates the discrimination threshold ⟨*d*⟩ = 2. (B–D) Shown are numerical simulations with *N*_L_ = 512, *μ* = *σ* = 1, and *λ* = 1 as well as *s* = 16 in (B,C) and *s* = 10 in (D).

The effect of varying receptor sensitivities can be studied in our model of primacy coding by discussing more general sensitivities matrices. We consider 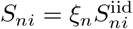, where each receptor type can have a different sensitivity factor *ξ_n_*, which modulates the uniform sensitivity matrix 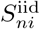 where each entry is independently chosen from the same log-normal distribution. The case of homogeneous sensitivities that we discusses so far thus corresponds to *ξ_n_* = 1.

To investigate the effect of heterogeneous sensitivities, we start by varying the sensitivity factor of one receptor type while keeping all others untouched, i. e., we change *ξ*_1_ while keeping *ξ_n_* = 1 for *n ≥* 2. There are three simple limits that we can discuss immediately. For *ξ*_1_ = 0, the first receptor type will never become active, the array behaves as if this type was not present, and the transmitted information is approximately *I*_max_(*N*_C_*, N*_R_ − 1). This value is lower than the maximally transmitted information *I*_max_(*N*_C_*, N*_R_) reached for the symmetric case *ξ*_1_ = 1. However, the associated information loss ∆*I* = *I*_max_(*N*_C_*, N*_R_) *− I*_max_(*N*_C_*, N*_R_ − 1) ≈ *N*_C_*/*(*N*_R_ ln 2) is relatively small in large receptor arrays (*N*_R_ ≫ *N*_C_); see Fig. 6B. Conversely, the transmitted information can be affected much more severely if the sensitivity of the first receptor type is increased beyond *ξ*_1_ = 1 and the receptors will thus be active more often than the others. In the extreme case of *ξ*_1_ → ∞, the first receptor type will always be active and thus not contribute any information. Since this receptor type would always be part of the primacy set, the information transmitted by the remaining receptor types is approximately *I*_max_(*N*_C_ − 1*, N*_R_ − 1), which is smaller than *I*_max_(*N*_C_*, N*_R_ 1) in the typical case *N*_R_ ≫ *N*_C_. Consequently, an overly active receptor type can be worse than not having this type at all under primacy coding.

The fact that overly sensitive receptors are detrimental to the transmitted information is also visible in numerical simulations. Fig. 6B shows ensemble averages of the information *I* transmitted by receptor arrays as a function of the sensitivity factor *ξ*_1_. As qualitatively argued above, *I* is maximal for *ξ*_1_ = 1 and it is slightly lower for smaller *ξ*_1_ since the receptor type is active less often. In contrast, for *ξ*_1_ > 1, *I* decreases dramatically and falls below the value of *ξ*_1_ = 0 for *ξ*_1_ ≳ 1.5. These data suggest that it would be better to remove receptor types that exhibit a 50 % higher sensitivity than the other types.

To see whether overly sensitive receptor types are also detrimental when all types have varying sensitivities, we next considering sensitivity factors *ξ_n_* distributed according to a lognormal distribution. Numerical results shown in Fig. 6C indicate that the transmitted information indeed decreases with increasing variance var(*ξ_n_*) of the sensitivity factors. In fact, a variation of var(*ξ_n_*)*/* ⟨*ξ_n_*⟩^2^ = 0.5 already implies a reduction of the transmitted information by almost 50 % for small concentration variations *σ/μ* = 1. If the odor concentrations vary more, the information degradation is less severe, but the same trend is visible. Interestingly, rescaling the information by the maximal information *I*_max_ given in Eq. (5) collapses the curves for all dimensions *N*_C_ and *N*_R_, suggesting that this analysis also holds for realistic receptor repertoire sizes. Note that the reduced transmitted information also implies poorer odor discrimination performance; see Fig. 6D. Taken together, this provides a strong selective pressure to limit the variability of the receptor sensitivities so overly sensitive receptors do not dominate the whole array.

## Discussion

We analyzed a simple model of primacy coding, where odors are identified by the *N*_C_ strongest responding receptor types. This primacy coding provides a sparse representation of the odor identity that is independent of the odor intensity. We showed using numerical simulations and a statistical model that the primacy dimension *N*_C_ strongly affects the transmitted information and the discriminability of odors. However, we showed that typical olfactory discrimination tasks can be carried out with performances close to experimentally measured ones for small *N*_C_ ≲ 10 already. Conversely, the number *N*_R_ of receptor types does not strongly affect the coding capacity and the discriminability of similar odors, in accordance with lesion experiments. Interestingly, our model even indicates that lowering *N*_R_ can improve the identification of a target ligand in a background.

Our model predicts that receptors need to respond with similar frequencies to incoming odors to be useful. This is because receptor types that are overly sensitive and respond strongly to many odors could dominate the response of other types and thus degrade the total information. In fact, having a receptor type that is 50 % more sensitive than others, and thus responds about three times as often, can lead to less transmitted information than when this type is absent. This observation is related to the primacy hull discussed in [37], which also predicts strong restrictions on the receptor sensitivities stemming from primacy coding. Various strategies could play a role in keeping the activity of the receptor types similar [46]: On timescales as short as a single sniff, the inhibition strength could be adjusted to regulate the relative importance of receptor excitations [47]. On longer timescales of several weeks, there are changes of the receptor copy number that directly affect the sensitivity of the glomeruli [48–50] and the processing neurons in the olfactory bulb [51, 52]. Receptor copy number adaptations influence the signal-to-noise ratio at the receptor level, so the copy number could be increased to improve the detection of frequently appearing odors [53]. In contrast, we predict a decrease of the copy number of overly sensitive receptor types that respond often. Combining the two alternatives, receptor copy numbers could be controlled such that noise is suppressed sufficiently while ensuring that single receptor types do not dominate the array. Finally, receptor sensitivities can also be adjusted by genetic modifications on evolutionary timescales [54, 55]. Moreover, direct feedback from higher regions of the brain could modify the processing of olfactory signals, e. g., in response to the behavioral state [7]. Although our work shows that the activities of the receptors need to be balanced, the actual distribution of the sensitivities matters much less. For instance, log-uniform distributions, which have been suggested to describe realistic receptor arrays [40, 56], lead to similar odor discriminability as log-normally distributed sensitivities; see Fig. S1.

Our results raise the question why mice have 20 times as many receptor types than flies, although the transmitted information under primacy coding is only increased by a factor of 2; see Eq. (5). The apparent usefulness of large receptor repertoires hints at roles of the olfactory system beyond transmitting the maximal information and discriminating average odors. For instance, having many receptor types might help to hardwire innate olfactory behavior when receptors are narrowly tuned to odors. In this case, our model would only apply to the fraction of the receptor types that are broadly tuned and are not connected to innate behavior. Alternatively, having many receptor types might be advantageous to discriminate very similar odor mixtures, to cover a larger dynamic range in concentrations of individual ligands, or to allow for a larger variation in average sensitivities, enabling quick adaptation to new environments. Finally, biophysical constraints of the receptor structure might imply that many receptors are required to cover a large part of chemical space.

Our model of primacy coding is very similar to our previous model of normalized receptor responses [21], which also exhibits concentration-invariant representations and predicts similar evolutionary pressure on the receptor sensitivities. In that case however, the mean activity decreases with larger mixture sizes, leading to diminishing discriminability of large mixtures caused by the constant inhibition strength [21]. Conversely, primacy coding can be interpreted as normalization with an inhibition strength that depends on the non-dimensional width of the concentration distribution; see Methods and Models. Primacy coding is thus an example for global inhibition with instantaneous adaptation, which displays better performance than a simple fixed threshold. Note that both models can detect targets in background odors, while this task is almost impossible without concentration invariance; see Fig. S2. Concentration invariance, here achieved by global inhibition, is therefore paramount for discriminating odors at various intensities. Taken together, our model suggests that primacy coding is superior at discriminating odors (Fig. 4C) while at the same time transmitting less information (Fig. 2A) compared to alternative models [21, 33, 40]. This implies that the information is more useful, which potentially allows for simpler processing downstream.

We discussed the simplest version of primacy coding with a minimal receptor model and a constant primacy dimension *N*_C_ implemented by a hard threshold. This model neglects the complex interactions of ligands at the olfactory receptors, which can affect perception [57]. In particular, antagonistic effects can already provide some normalization at the level of receptors [58]. Generally, it is likely that many mechanisms contribute to the overall normalization of the receptor response [59]. A more realistic model of primacy coding might also consider a softer threshold, where receptor types with larger excitation are given higher weight in the downstream interpretation, which is related to rank coding [22]. In this case, information from fewer glomeruli might be sufficient to identify odors, since the rank carries additional information; see Fig. 2A. Realistic olfactory systems could also use a timing code, taking into account more and more receptor types (with decreasing excitation) until an odor is identified confidently. Such a system could explain that the response dynamics in experiment depend on the task [60, 61]. Generally, a better understanding of the temporal structure of the olfactory code [8, 62–66] might allow to derive more detailed models. These could rely on attractor dynamics that are guided by the excitations and thus respond stronger to the early and large excitations [67, 68].

## Methods and Models

### Numerical simulations

All numerical simulations are based on ensemble averages over odors ***c*** and sensitivity matrices *S_ni_*. The elements of *S_ni_* are drawn independently from a log-normal distribution with var(*S_ni_*)*/S*̄^2^ = 1.72 corresponding to *λ* = 1. Odors ***c*** are chosen by first determining which of the *N*_L_ ligands are present using a Bernoulli distribution with probability *p* = *s/N*_L_ and then drawing their concentration from a log-normal distribution with mean *μ* and standard deviation *σ*. The primacy set ***a*** corresponding to ***c*** is given by the *N*_C_ receptors with the highest excitation calculated from Eq. (1). Statistics of ***a*** and the transmitted information *I* given by Eq. (4) are determined by repeating this procedure 10^5^ and 10^7^ times, respectively.

### Statistical model

The statistics of the output ***a*** given by Eqs. (1)–(3) can be estimated using ensemble averages of sensitivity matrices for different odors ***c***, similar to our treatment presented in [21] and [33]. In particular, Eq. (1) implies that the excitations *e_n_* are well approximated by a log-normal distribution with mean 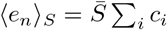 and variance and 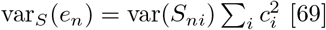) whereas correlations are negligible [21]. The probability that the excitation *e_n_* exceeds the threshold *γ* and the associated receptor type is part of the primacy set reads

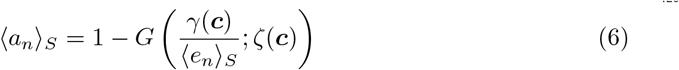

with

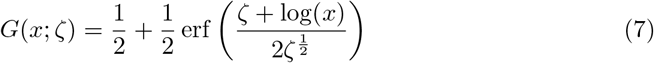

being the cumulative density function of a log-normal distribution with ⟨*x*⟩ = 1 and var(*x*) = exp(2*ζ*) − 1. The width of the distribution is determined by the positive parameter 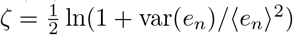), which reads

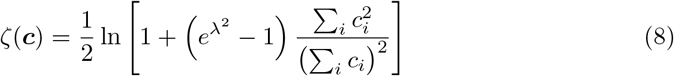

for an ensemble average over sensitivities. Note that *ζ* is concentration-invariant, since it does not change when the concentration vector ***c*** is multiplied by a constant factor. In the simple case of ligands that are distributed according to *P*_env_(***c***), we find 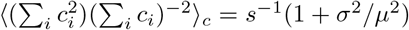. Consequently, the distribution width *ζ* is large for broadly distributed sensitivities (large *λ*), few ligands in an odor (small *s*), and wide concentration distributions (large *σ/μ*).

The constraint Eq. (3) implies ⟨*a_n_*⟩ = *N*_C_*/N*_R_, so that the mean threshold reads

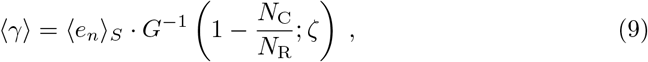

where *G^−^*^1^ is the inverse function of *G* defined in Eq. (7). Using this expression as an estimate for *γ* in Eq. (6) results in concentration-invariant activities *a_n_*, since ⟨*γ*⟩ is proportional to the excitation ⟨*e_n_*⟩. This situation is comparable to simple normalized representations resulting from the threshold *γ* = *α* ⟨*e_n_*⟩, where *α* is a constant inhibition strength [21]. In fact, primacy coding can be interpreted as global inhibition with an inhibition threshold depending on the width of the excitation distribution, *α* = *G^−^*^1^(1 *− N*_C_*N*_R_^−1^; *ζ*).

#### Inter-excitation intervals

The expected difference between excitations corresponding to a given odor ***c*** can be studied using order statistics, where excitations are re-indexed such that they are ordered, 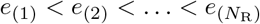. For simplicity, we consider the case where the excitations *e_n_* are distributed identically when considering all odors according to *P*_env_(***c***). Denoting the cumulative distribution function of the excitations by 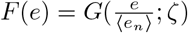 and the associated probability density function by *f* (*e*), the probability density function associated with the excitation *e*_(*n*)_ at rank *n* reads [70]

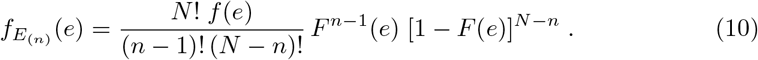

The joint distribution of *E*_(*n*)_ and *E*_(*m*)_, 1 *≤ n < m ≤ N*, reads [70]

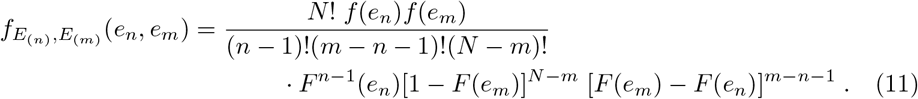

Consequently, the distribution of the difference ∆*e* = *e*_(*n*)_ − *e*_(*n−*1)_ of consecutive excitations is

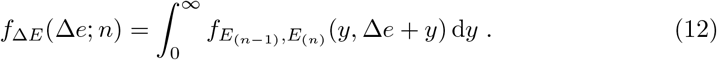

Hence, the expected difference ⟨∆*e*⟩ = ∫ *x f*_∆*E*_(*x*; *N*_R_ *N*_C_ 1) d*x* between the strongest excited inactive receptor type and the weakest active receptor type can be evaluated.

#### Distances between primacy set

The expected number ⟨*d*⟩ of changes in the primacy set ***a*** when a target odor ***c***^t^ is added to some background ***c***^b^ reads

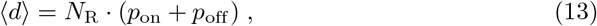

where *p*_on_ is the probability that a receptor type that was inactive for ***c***^b^ is turned on by the perturbation ***c***^t^ and *p*_off_ is the probability that a receptor type that was active is turned off. Both probabilities depend on the excitation thresholds *γ*^(1)^ and *γ*^(2)^ associated with the odors ***c***^b^ and ***c***^b^ + ***c***^t^, respectively, which can be estimated from Eq. (9) using the respective excitation statistics. With this, *p*_on_ follows from the probability that the excitation was at the value *x* below *γ*^(1)^ and the additional excitation by the target brings the total excitation above *γ*^(2)^,

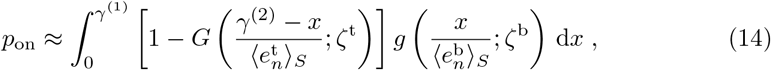

where *g*(*e*;*ζ*) is the probability density function associated with *G*(*e*;*ζ*) given in Eq. (7). Here, 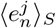 and *ζ^j^* describe the excitation statistics of the target (*j* = t) and the background (*j* = b). Similarly, we obtain

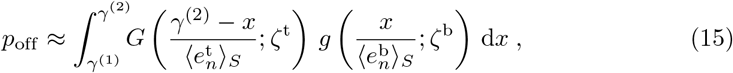

so we can use Eq. (13) to calculate the expected Hamming distance ⟨*d*⟩. Note that *γ*^(1)^ and *γ*^(2)^ depend on *N*_R_, so the distance ⟨*d*⟩ does thus not scale trivially with *N*_R_, in contrast to the case of normalized representations [21].

We use Eqs. (13)–(15) to calculated ⟨*d*⟩ when a target ligand with concentration *c*_t_ is added to a background ligand at concentration *c*_b_. The associated statistics of the excitations obey

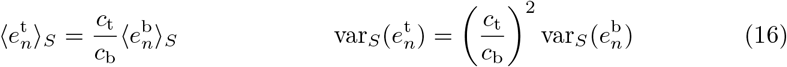

and 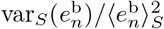 follows from chosen values of *σ/μ* and *λ*. Similarly, when a ligand with concentration *c* is added to a mixture of *s* ligands, all at concentration *c*, we have

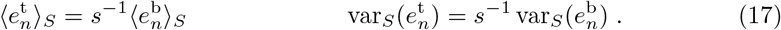

The third case of correlated odors that we discuss in the main text concerns two odor mixtures of equal size *s* sharing *s*_B_ of the ligands. In this case, the excitation threshold *γ* is the same for both odors and we can express the probability *p*_xor_ that a receptor type is excited by one mixture but not the other as

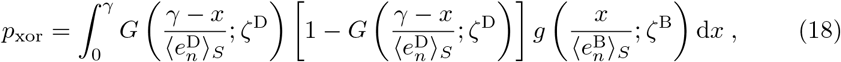

where the statistics 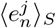 and *ζ^j^* need to be evaluated for the excitations associated with the *s*_B_ ligands that are the same (*j* = B) and the *s − s*_B_ ligands that are different (*j* = D) between the two mixtures. Taken together, the expected distance reads ⟨*d*⟩ = 2*N*_R_*p*_xor_ and we recover ⟨*d*⟩= *d_∗_* for unrelated mixtures (*s*_B_ = 0) and ⟨*d*⟩= 0 for identical mixtures (*s*_B_ = *s*).

#### Information transmitted by diverse receptors

In the case where the primacy sets ***a*** can be partitioned into *N_M_* groups with all elements within a group appearing with the same probability, we can write the information *I* given by Eq. (4) as

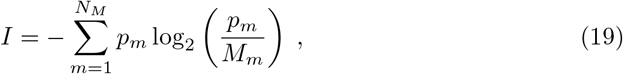

where *M_m_* is the number of elements within group *m* and *p_m_* is the probability that group *m* appears in the output, such that 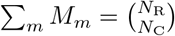 and 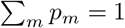. In the simple case of one receptor type with deviating statistics, we have *N_M_* = 2 with

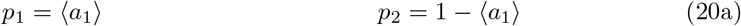

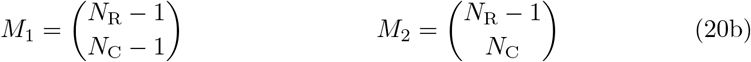

while the remaining activities are ⟨*a_n_*⟩= (*N*_C_ − ⟨*a*_1_⟩)*/*(*N*_R_ − 1) for *n ≥* 2 to obey Eq. (3). For *p*_1_ = 0, Eq. (19) reduces to *I* = *I*_max_(*N*_C_*, N*_R_ − 1), whereas the maximum *I* = *I*_max_(*N*_C_*, N*_R_) is reached for *p*_1_ = *N*_C_*/N*_R_. The information decreases for larger *p*_1_ and eventually reaches values lower than *I*_max_(*N*_C_*, N*_R_ 1) when 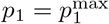. For 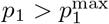, it would thus be advantageous to remove this receptor type. Using 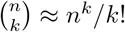 and expanding Eq. (19) around *p*_1_ = *eN*_R_*/*(*N*_R_ − 1), we find

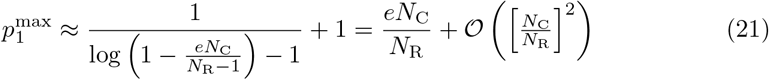

in the limit *N*_R_ ≫ *N*_C_ of large repertoires, so 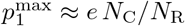.

## Supporting information

**S1 Fig. Log-uniform distributed sensitivities behave similar to log-normal distributed ones under primacy coding.** Analysis of odor discrimination of primacy coding with log-uniform distributed sensitivities.

**S2 Fig. Target detection fails without concentration normalization.** Analysis of odor discrimination of a binary encoding with log-normal and log-uniform sensitivities.

## Acknowledgments

This work was funded by the Simons Foundation and the German Science Foundation through ZW 222/1-1.

---

**Fig S1.**
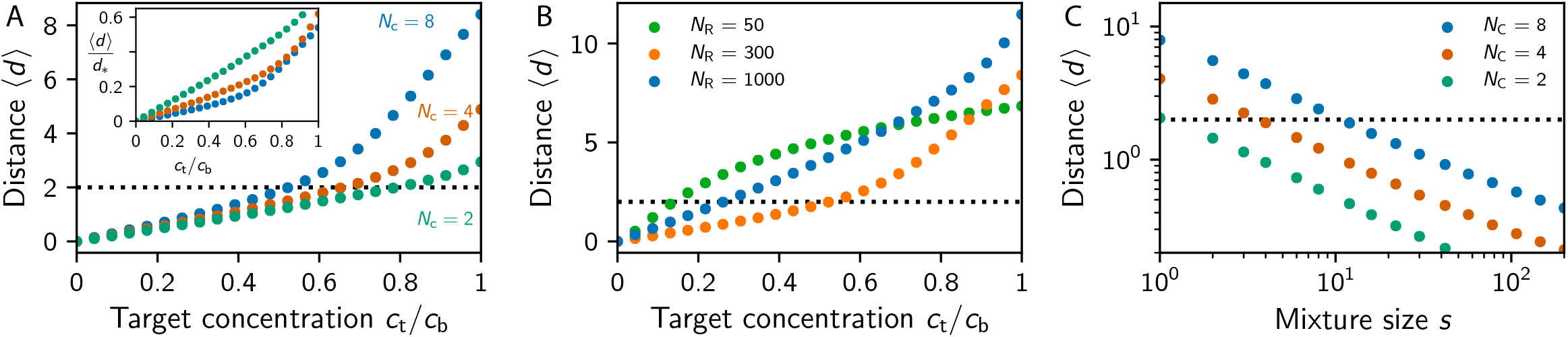
Log-uniform distributed sensitivities behave similar to log-normal distributed ones under primacy coding. (A-B) Mean change ⟨*d*⟩in the primacy set ***a*** when a target ligand at concentration *c*_t_ is added to a background ligand at concentration *c*_b_ as a function of *c*_t_*/c*_b_ for (A) various *N*_C_ at *N*_R_ = 300 (inset: same data rescaled by *d_∗_*) and (B) various *N*_R_ at *N*_C_ = 8. (C) Mean change ⟨*d*⟩in the primacy set when a ligand is added to a mixture with *s* ligands as a function of *s* for various *N*_C_ and *N*_R_ = 300. (A–C) The dotted line indicates the discrimination threshold ⟨*d*⟩= 2. Shown are numerical simulations (dots; sample size: 10^5^) for *N*_L_ = 512, *σ/μ* = 0, and var(*S_ni_*)*/S*̄^2^= 7, so the log-uniform distributed sensitivities span 7 orders of magnitude. Note that the three panels are similar to Fig. 3A, 3B and 4A, respectively, implying that log-uniform and log-normal distributed *S_ni_* behave similarly.

**Fig S2.**
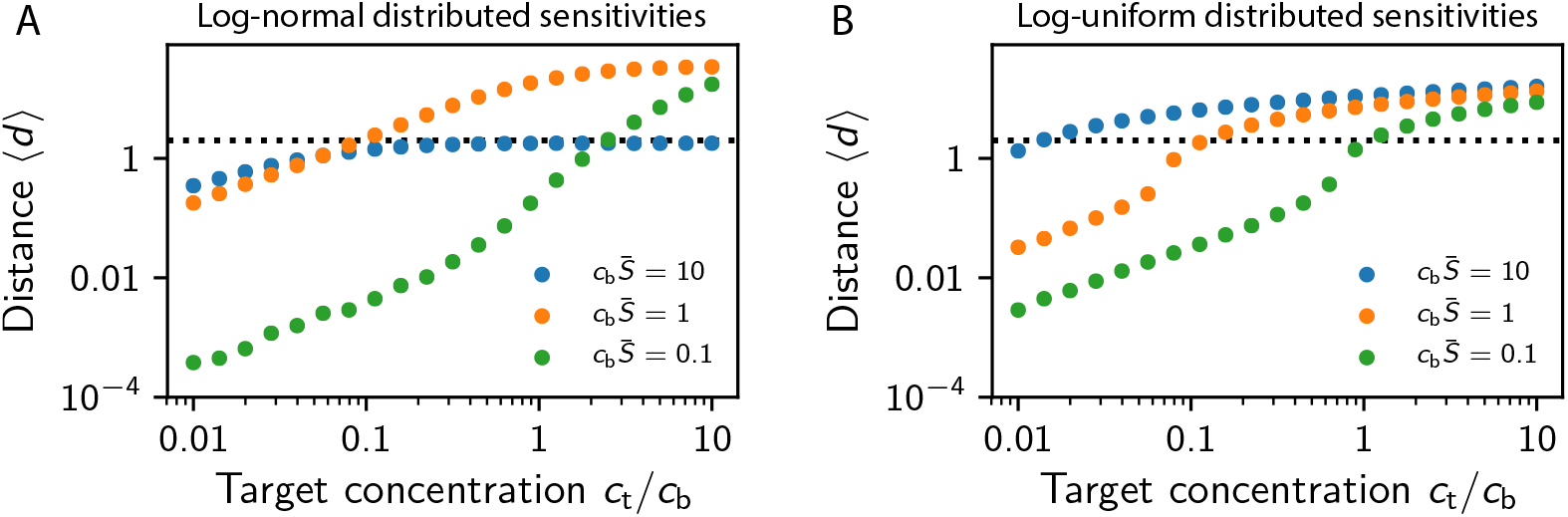
Target detection fails without concentration normalization. Mean change ⟨*d*⟩ in the activity ***a*** associated with binary coding (*a_n_* = 1 if and only if *e_n_ >* 1) when a target ligand at concentration *c*_t_ is added to a background ligand at concentration *c*_b_ as a function of *c*_t_*/c*_b_ for various concentrations *c*_b_. Shown are numerical simulations (dots; sample size: 10^5^) for *N*_R_ = *N*_L_ = 50 and *σ/μ* = 0. (A) Log-normal distributed *S_ni_* with var(*S_ni_*)*/S*̄^2^ = 1.72 discussed in [33]. (B) Log-uniform distributed *S_ni_* with var(*S_ni_*)*/S*̄^2^ = 7 discussed in [40].

## References

1. Malnic B, Hirono J, Sato T, Buck LB. Combinatorial receptor codes for odors. Cell. 1999;96(5):713–723.

2. Barnes DC, Hofacer RD, Zaman AR, Rennaker RL, Wilson DA. Olfactory perceptual stability and discrimination. Nat Neurosci. 2008;11(12):1378–80. doi:10.1038/nn.2217.

3. Mathis A, Rokni D, Kapoor V, Bethge M, Murthy VN. Reading out olfactory receptors: feedforward circuits detect odors in mixtures without demixing. Neuron. 2016;91(5):1110–1123.

4. Tabor R, Yaksi E, Weislogel JM, Friedrich RW. Processing of odor mixtures in the zebrafish olfactory bulb. J Neurosci. 2004;24(29):6611–20. doi:10.1523/JNEUROSCI.1834-04.2004.

5. Bolding KA, Franks KM. Complementary codes for odor identity and intensity in olfactory cortex. Elife. 2017;6. doi:10.7554/eLife.22630.

6. Roland B, Deneux T, Franks KM, Bathellier B, Fleischmann A. Odor identity coding by distributed ensembles of neurons in the mouse olfactory cortex. Elife. 2017;6. doi:10.7554/eLife.26337.

7. Wilson RI. Early olfactory processing in Drosophila: mechanisms and principles. Annu Rev Neurosci. 2013;36:217–41. doi:10.1146/annurev-neuro-062111-150533.

8. Uchida N, Poo C, Haddad R. Coding and transformations in the olfactory system. Annu Rev Neurosci. 2014;37:363–85. doi:10.1146/annurev-neuro-071013-013941.

9. Silva Teixeira CS, Cerqueira NMFSA, Silva Ferreira AC. Unravelling the Olfactory Sense: From the Gene to Odor Perception. Chem Senses. 2016;41(2):105–21. doi:10.1093/chemse/bjv075.

10. Yokoi M, Mori K, Nakanishi S. Refinement of odor molecule tuning by dendrodendritic synaptic inhibition in the olfactory bulb. Proc Natl Acad Sci USA. 1995;92(8):3371–5.

11. Berck ME, Khandelwal A, Claus L, Hernandez-Nunez L, Si G, Tabone CJ, et al. The wiring diagram of a glomerular olfactory system. Elife. 2016;5. doi:10.7554/eLife.14859.

12. Li Z. A model of olfactory adaptation and sensitivity enhancement in the olfactory bulb. Biol Cybern. 1990;62(4):349–61.

13. Li Z. Modeling the sensory computations of the olfactory bulb. In: Models of neural networks. Springer; 1994. p. 221–251.

14. Linster C, Hasselmo M. Modulation of inhibition in a model of olfactory bulb reduces overlap in the neural representation of olfactory stimuli. Behavioural brain research. 1997;84(1):117–127.

15. Cleland TA, Sethupathy P. Non-topographical contrast enhancement in the olfactory bulb. BMC Neurosci. 2006;7:7. doi:10.1186/1471-2202-7-7.

16. Olsen SR, Bhandawat V, Wilson RI. Divisive normalization in olfactory population codes. Neuron. 2010;66(2):287–99. doi:10.1016/j.neuron.2010.04.009.

17. Zhang D, Li Y, Wu S. Concentration-invariant odor representation in the olfactory system by presynaptic inhibition. Comput Math Methods Med. 2013;2013:507143. doi:10.1155/2013/507143.

18. Roland B, Jordan R, Sosulski DL, Diodato A, Fukunaga I, Wickersham I, et al. Massive normalization of olfactory bulb output in mice with a ‘monoclonal nose’. Elife. 2016;5. doi:10.7554/eLife.16335.

19. Laurent G. A systems perspective on early olfactory coding. Science. 1999;286(5440):723–8.

20. Cleland TA. Early transformations in odor representation. Trends Neurosci. 2010;33(3):130–139. doi:10.1016/j.tins.2009.12.004.

21. Zwicker D. Normalized Neural Representations of Complex Odors. PLoS One. 2016;11(11):e0166456. doi:10.1371/journal.pone.0166456.

22. Junek S, Kludt E, Wolf F, Schild D. Olfactory coding with patterns of response latencies. Neuron. 2010;67(5):872–84. doi:10.1016/j.neuron.2010.08.005.

23. Wilson CD, Serrano GO, Koulakov AA, Rinberg D. A primacy code for odor identity. Nat Commun. 2017;8(1):1477. doi:10.1038/s41467-017-01432-4.

24. Wesson DW, Carey RM, Verhagen JV, Wachowiak M. Rapid encoding and perception of novel odors in the rat. PLoS Biol. 2008;6(4):e82. doi:10.1371/journal.pbio.0060082.

25. Kepple D, Giaffar H, Rinberg D, Koulakov A. Deconstructing Odorant Identity via Primacy in Dual Networks. arXiv preprint arXiv:160902202. 2016;.

26. Dunkel M, Schmidt U, Struck S, Berger L, Gruening B, Hossbach J, et al. SuperScent–a database of flavors and scents. Nucleic Acids Res. 2009;37(Database issue):D291–4. doi:10.1093/nar/gkn695.

27. Touhara K, Vosshall LB. Sensing odorants and pheromones with chemosensory receptors. Annu Rev Physiol. 2009;71:307–32. doi:10.1146/annurev.physiol.010908.163209.

28. Wright GA, Thomson MG. Odor Perception and the Variability in Natural Odor Scenes. In: Romeo J, editor. Integrative Plant Biochemistry. vol. 39 of Recent Advances in Phytochemistry. Elsevier; 2005. p. 191–226.

29. Verbeurgt C, Wilkin F, Tarabichi M, Gregoire F, Dumont JE, Chatelain P. Profiling of olfactory receptor gene expression in whole human olfactory mucosa. PLOS ONE. 2014;9(5):e96333. doi:10.1371/journal.pone.0096333.

30. Niimura Y. Olfactory receptor multigene family in vertebrates: from the viewpoint of evolutionary genomics. Curr Genomics. 2012;13(2):103–14. doi:10.2174/138920212799860706.

31. Su CY, Menuz K, Carlson JR. Olfactory perception: receptors, cells, and circuits. Cell. 2009;139(1):45–59. doi:10.1016/j.cell.2009.09.015.

32. Silbering AF, Galizia CG. Processing of odor mixtures in the Drosophila antennal lobe reveals both global inhibition and glomerulus-specific interactions. J Neurosci. 2007;27(44):11966–77. doi:10.1523/JNEUROSCI.3099-07.2007.

33. Zwicker D, Murugan A, Brenner MP. Receptor arrays optimized for natural odor statistics. Proc Natl Acad Sci USA. 2016;113(20):5570–75. doi:10.1073/pnas.1600357113.

34. Mainland JD, Li YR, Zhou T, Liu WLL, Matsunami H. Human olfactory receptor responses to odorants. Sci Data. 2015;2:150002. doi:10.1038/sdata.2015.2.

35. Münch D, Galizia CG. DoOR 2.0 - Comprehensive Mapping of Drosophila melanogaster Odorant Responses. Sci Rep. 2016;6:21841. doi:10.1038/srep21841.

36. Gao XJ, Clandinin TR, Luo L. Extremely sparse olfactory inputs are sufficient to mediate innate aversion in Drosophila. PLoS One. 2015;10(4):e0125986. doi:10.1371/journal.pone.0125986.

37. Giaffar H, Rinberg D, Koulakov AA. Primacy model and the evolution of the olfactory receptor repertoire. bioRxiv. 2018; p. 255661.

38. Jefferis GS, Marin EC, Stocker RF, Luo L. Target neuron prespecification in the olfactory map of Drosophila. Nature. 2001;414(6860):204–8. doi:10.1038/35102574.

39. Cleland TA, Linster C. On-Center/Inhibitory-Surround Decorrelation via Intraglomerular Inhibition in the Olfactory Bulb Glomerular Layer. Front Integr Neurosci. 2012;6:5. doi:10.3389/fnint.2012.00005.

40. Koulakov A, Gelperin A, Rinberg D. Olfactory coding with all-or-nothing glomeruli. J Neurophysiol. 2007;98(6):3134–3142.

41. Slotnick B, Bisulco S. Detection and discrimination of carvone enantiomers in rats with olfactory bulb lesions. Neuroscience. 2003;121(2):451–7.

42. Bisulco S, Slotnick B. Olfactory discrimination of short chain fatty acids in rats with large bilateral lesions of the olfactory bulbs. Chem Senses. 2003;28(5):361–70.

43. Jinks A, Laing DG. A limit in the processing of components in odour mixtures. Perception. 1999;28(3):395–404.

44. Rokni D, Hemmelder V, Kapoor V, Murthy VN. An olfactory cocktail party: figure-ground segregation of odorants in rodents. Nat Neurosci. 2014;17(9):1225–32. doi:10.1038/nn.3775.

45. Bushdid C, Magnasco M, Vosshall L, Keller A. Humans can discriminate more than 1 trillion olfactory stimuli. Science. 2014;343(6177):1370–1372.

46. Zufall F, Leinders-Zufall T. The cellular and molecular basis of odor adaptation. Chem Senses. 2000;25(4):473–81.

47. Hong EJ, Wilson RI. Simultaneous encoding of odors by channels with diverse sensitivity to inhibition. Neuron. 2015;85(3):573–89. doi:10.1016/j.neuron.2014.12.040.

48. Lazarini F, Lledo PM. Is adult neurogenesis essential for olfaction? Trends Neurosci. 2011;34(1):20–30. doi:10.1016/j.tins.2010.09.006.

49. Yu CR, Wu Y. Regeneration and rewiring of rodent olfactory sensory neurons. Exp Neurol. 2016;doi:10.1016/j.expneurol.2016.06.001.

50. Ibarra-Soria X, Nakahara TS, Lilue J, Jiang Y, Trimmer C, Souza MA, et al. Variation in olfactory neuron repertoires is genetically controlled and environmentally modulated. Elife. 2017;6. doi:10.7554/eLife.21476.

51. Mouret A, Lepousez G, Gras J, Gabellec MM, Lledo PM. Turnover of newborn olfactory bulb neurons optimizes olfaction. J Neurosci. 2009;29(39):12302–14. doi:10.1523/JNEUROSCI.3383-09.2009.

52. Liu A, Savya S, Urban NN. Early Odorant Exposure Increases the Number of Mitral and Tufted Cells Associated with a Single Glomerulus. J Neurosci. 2016;36(46):11646–11653. doi:10.1523/JNEUROSCI.0654-16.2016.

53. Tesileanu T, Cocco S, Monasson R, Balasubramanian V. Environmental adaptation of olfactory receptor distributions. arXiv preprint arXiv:180109300. 2018;.

54. Yu Y, Claire A, Ni MJ, Adipietro KA, Golebiowski J, Matsunami H, et al. Responsiveness of G protein-coupled odorant receptors is partially attributed to the activation mechanism. Proc Natl Acad Sci USA. 2015;112(48):14966–14971.

55. Bear DM, Lassance JM, Hoekstra HE, Datta SR. The Evolving Neural and Genetic Architecture of Vertebrate Olfaction. Curr Biol. 2016;26(20):R1039–R1049. doi:10.1016/j.cub.2016.09.011.

56. Hopfield J. Odor space and olfactory processing: collective algorithms and neural implementation. Proc Natl Acad Sci USA. 1999;96(22):12506–12511.

57. Saraiva LR, Kondoh K, Ye X, Yoon KH, Hernandez M, Buck LB. Combinatorial effects of odorants on mouse behavior. Proc Natl Acad Sci USA. 2016;doi:10.1073/pnas.1605973113.

58. Reddy G, Zak JD, Vergassola M, Murthy VN. Antagonism in olfactory receptor neurons and its implications for the perception of odor mixtures. Elife. 2018;7. doi:10.7554/eLife.34958.

59. Cleland TA, Chen SYT, Hozer KW, Ukatu HN, Wong KJ, Zheng F. Sequential mechanisms underlying concentration invariance in biological olfaction. Front Neuroeng. 2011;4:21. doi:10.3389/fneng.2011.00021.

60. Schaefer AT, Margrie TW. Psychophysical properties of odor processing can be quantitatively described by relative action potential latency patterns in mitral and tufted cells. Front Syst Neurosci. 2012;6:30. doi:10.3389/fnsys.2012.00030.

61. Frederick DE, Brown A, Tacopina S, Mehta N, Vujovic M, Brim E, et al. Task-Dependent Behavioral Dynamics Make the Case for Temporal Integration in Multiple Strategies during Odor Processing. J Neurosci. 2017;37(16):4416–4426. doi:10.1523/JNEUROSCI.1797-16.2017.

62. Egea-Weiss A, Kleineidam CJ, Szyszka P, et al. High Precision of Spike Timing across Olfactory Receptor Neurons Allows Rapid Odor Coding in Drosophila. iScience. 2018;4:76–83.

63. Gupta P, Albeanu DF, Bhalla US. Olfactory bulb coding of odors, mixtures and sniffs is a linear sum of odor time profiles. Nat Neurosci. 2015;18(2):272–81.

64. Blauvelt DG, Sato TF, Wienisch M, Murthy VN. Distinct spatiotemporal activity in principal neurons of the mouse olfactory bulb in anesthetized and awake states. Frontiers in neural circuits. 2013;7(46).

65. Spors H, Wachowiak M, Cohen LB, Friedrich RW. Temporal dynamics and latency patterns of receptor neuron input to the olfactory bulb. J Neurosci. 2006;26(4):1247–59. doi:10.1523/JNEUROSCI.3100-05.2006.

66. Hopfield JJ. Pattern recognition computation using action potential timing for stimulus representation. Nature. 1995;376(6535):33–6. doi:10.1038/376033a0.

67. Herman PA, Benjaminsson S, Lansner A. Odor recognition in an attractor network model of the mammalian olfactory cortex. In: Neural Networks (IJCNN), 2017 International Joint Conference on. IEEE; 2017. p. 3561–3568.

68. Grabska-Barwińska A, Barthelmé S, Beck J, Mainen ZF, Pouget A, Latham PE. A probabilistic approach to demixing odors. Nat Neurosci. 2016;doi:10.1038/nn.4444.

69. Fenton LF. The sum of log-normal probability distributions in scatter transmission systems. Communications Systems, IRE Transactions on. 1960;8(1):57–67.

70. David HA, Nagaraja HN. Order statistics. Wiley Online Library; 1970.

